# Draft genome assembly of the Woolly bottlebrush (*Greyia radlkoferi*) using Pacbio long-read sequencing technology

**DOI:** 10.64898/2026.05.26.727846

**Authors:** A.H. Molotsi, T. Masebe, L.T. Nesengani, S. Mdyogolo, T.S. Tshilate, R.M. Smith, N. Hlongwane, S. Hadebe, T.M. Mafokwane, N. Mapholi

**Affiliations:** Department of Agriculture and Animal Health, College of Agriculture and Environmental Sciences, University of South Africa; Department of Life and Consumer Sciences, College of Agriculture and Environmental Sciences, University of South Africa

## Abstract

The Woolly bottlebrush (*Greyia radlkoferi*) is an indigenous South African plant known for its ornamental appeal and potential medicinal uses. It naturally grows on rocky hillsides and grasslands and highly resilient to drought, temperature fluctuations, and nutrient-poor soils. Its flavonoid-rich compounds with anti-tyrosinase activity support its traditional use to treat skin pigmentation disorders in humans. Despite its outstanding ecological and biochemical characteristics, no reference genome is available for *Greyia radlkoferi*. Therefore, this study aimed to generate the first draft genome of the *Greyia Radlkofleri* using PacBio Sequel IIe HiFi long read sequencing. A total of 56.07 Gb HiFi data was generated, providing a total genome coverage of 270X. The assembled genome size was 206Mb, with the longest scaffold being 13.9 Mb. The assembly statistics yielded a scaffold and contig N50 of 10.1 Mb, and an L50 of 9, and with an overall GC content of 34.9 %. The genome scope profile set at kmer = 17 indicated that the genome is triploid. The genome annotation predicted 17,804 protein-coding genes and 17,804 transcripts with an average gene length of 3,116.03 bp. This is the first draft genome of its kind for the *Greyia* genus and provides a foundation for future studies aimed at elucidating the genetic basis of its environmental resilience and the biosynthetic pathways underlying its medicinal properties.

## Background and Summary

The Woolly bottlebrush (*Greyia radlkoferi*) (Figure 1) is one of the three species of the *Greyia* genus found in South Africa specifically, in Limpopo province^1^. The other species in the *Greyia* genus include the *G. flanaganii* Bolus and the *G. sutherlandii* which are found in the Eastern Cape and Kwa-Zulu Natal provinces, respectively. The *G. radlkoferi species* is a deciduous shrub which can grow up to 5 m in height. The plant is characterised by red flowers which open into a bottlebrush, with its leaves having a white woolly appearance on the abaxial surface. Greyia species are known to produce rich nectar during late winter and early spring which acts as a pollinator for bees, birds and butterflies^2^. The *Greyia* genus is considered a distant relative of *Melianthus* ^3^, with both belonging to the order of *Geraniales* in the eudicot clade. Phylogenetic relationships within the groups were resolved through DNA barcoding approaches based on ITS and *trnL-F* gene regions^3,4^ with *Greyia* classified within the *Francoaceae* Familys^4^.

**Figure 1.**
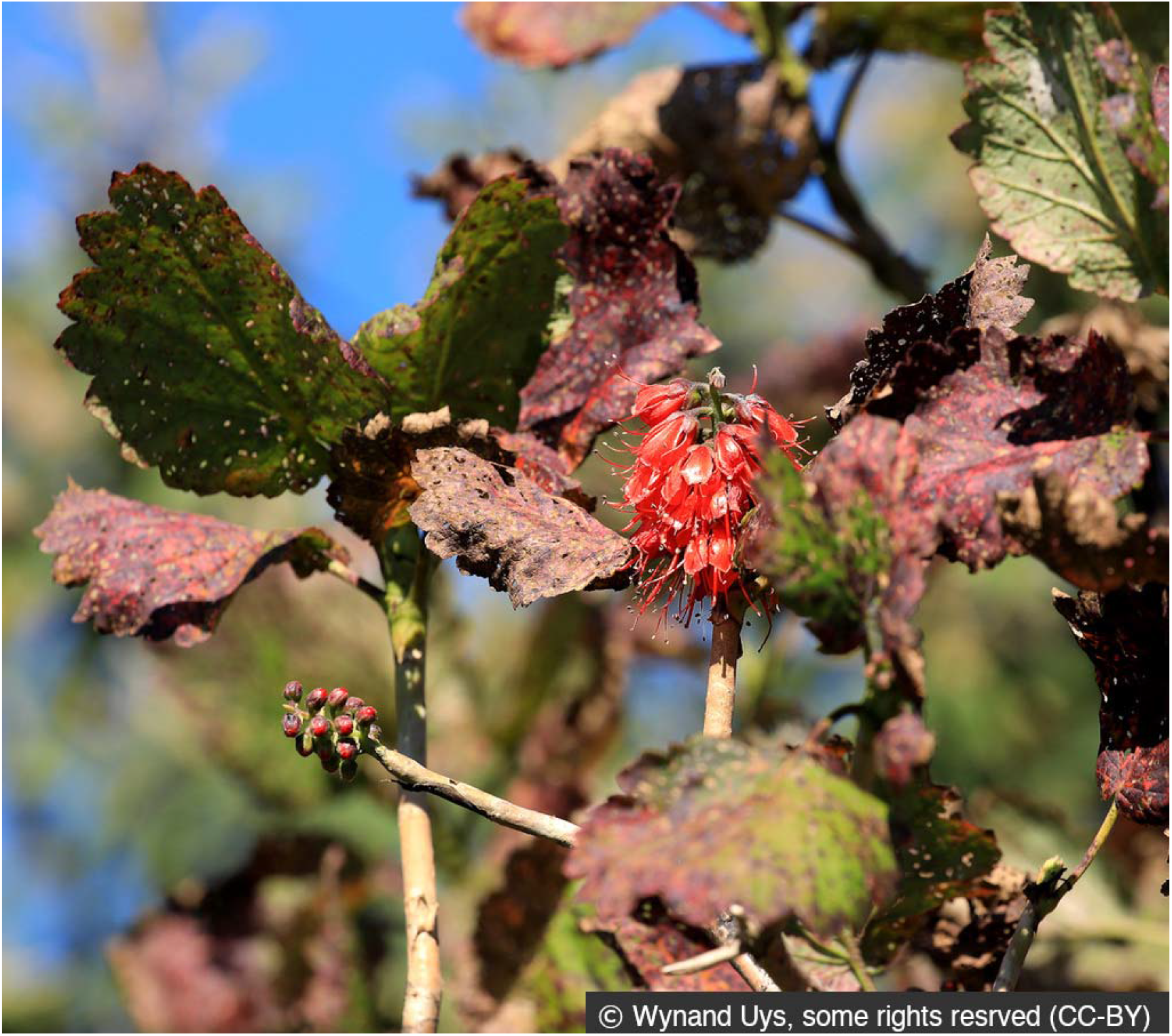
Woolly Bottlebrush in its natural habitat. Photo © Wynand Uys via iNaturalist (Observation # 221795082), licensed under CC-BY 4.0. Link: https://www.inaturalist.org/observations/221795082

In addition to its taxonomic and evolutionary significance, the plant is widely recognized for its medicinal properties, traditionally used to treat skin pigmentation. The key bioactive compound of *G. radlkoferi* is 2’,4’,6’ trihydroxydihydrochalcone, which has anti-tyrosinase activity known to inhibit the production of melanin^5^. Consequently, most studies on this species have focused on metabolomic analysis to characterise the bioactive compounds for medicinal purposes^6–9^; however, there have been no genomic reported for *G. radlkoferi* in South Africa.

The increasing use of *G. radlkoferi* as traditional medicine highlights the need for accurate species identification, characterization and authentication. However, current DNA barcoding approaches are limited to genetic characterisation and resolving species-level variation. Recent studies have emphasised the need for utilisation of SNP markers to provide species-specific authentication of *Greyia* extracts in herbal products. To identify species-specific SNP markers in *Greyia*, it is crucial to have the whole genome sequencing data for these species. Currently, three plant species within the order of Geraniales have been sequenced to contig level using Illumina short-read sequencing. These plants are *Melianthus villosus, Francoa sonchifolia and Viviania marifolia*. Therefore, this study aimed to generate a draft genome of *G. radlkoferi* and conduct ortholog analysis of *G. radlkoferi* with related plant species within the Eudicots clade.

## Methods

### Sample collection and sequencing

*G. radlkoferi* plant tissue samples were obtained from SANBI biobanking facilities in Pretoria. The samples were immediately placed on dry ice, transported to the laboratory, and stored at −80°C until further processing. Genomic DNA was extracted from the plant tissue using the Nanobind protocol for high-molecular-weight (HMW) DNA extraction. Thereafter, the SMRTBell® prep kit 3.0 was used to prepare the sequencing library. Sequencing was then performed using the PacBio Sequel IIe platform generating a total data output of 56.07 Gb, thus providing a total coverage of approximately 270X. Initial sequence quality control processing was performed using the SMRTlink Software v11.0 (PacBio) as well as FastQC v0.12.1^10^.

### Genome assembly pipeline

The genome assembly was carried out through the reproducible Vertebrate Genome Project (VGP) workflows on the European Galaxy platform^11^. These workflows begin with utilisation of GenomeScope2 Version 2.1.0^12^ for genome characterization and estimation of genome size from HiFi reads, which resulted in an k-mer based genome size of 235 bp for *G. radlkoferi* from k-mer (17) analysis (Figure 2). Additionally, smudgeplot v2.0.5^13^ analysis was also run to determine ploidy level of the genome (Figure 3).

**Figure 2.**
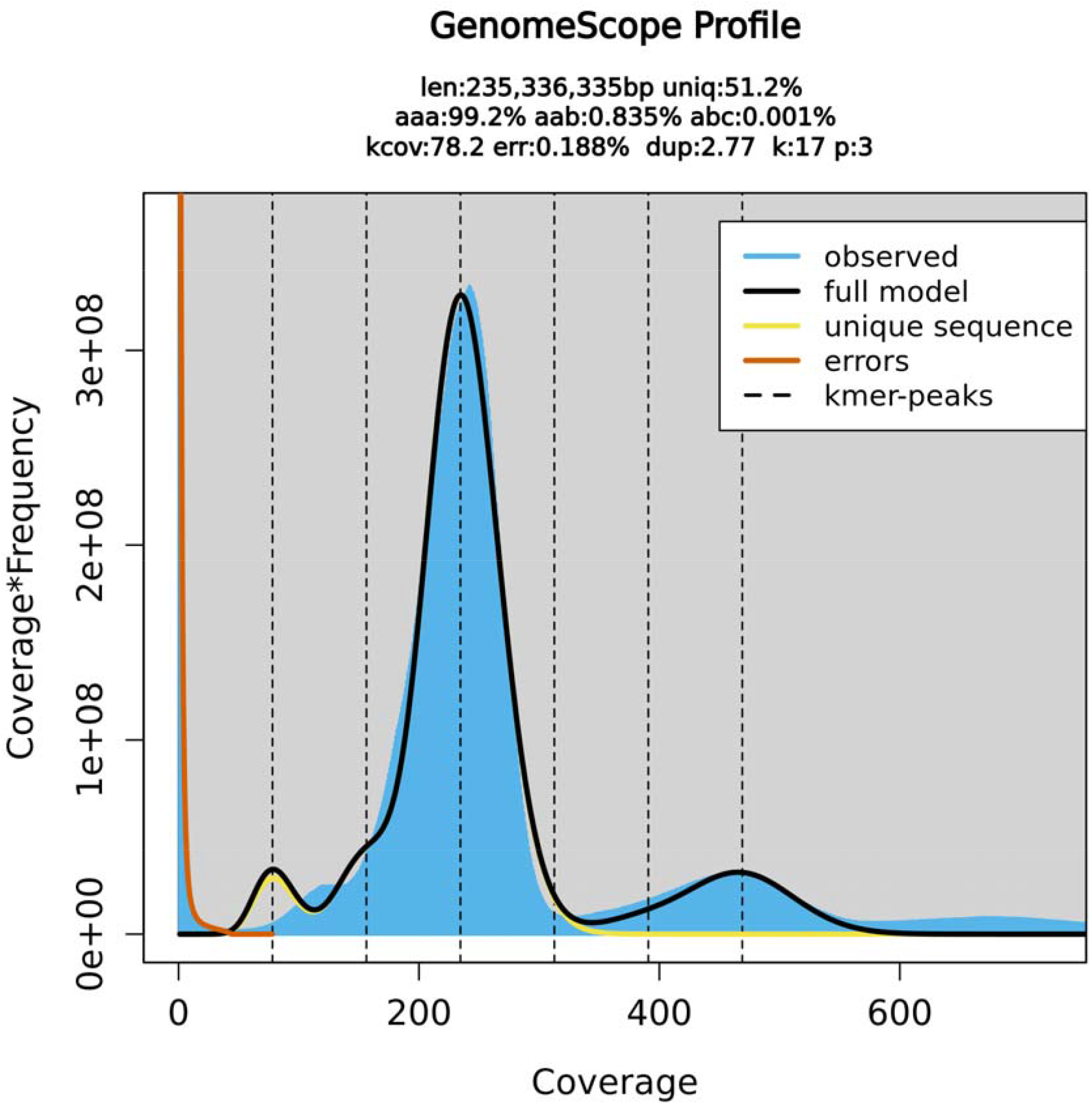
Genome scope profile for the *G. radlkoferi* genome. The figure describes the estimated genome size (len), the heterozygosity (ab), and homozygosity (aa), the number of duplicates (dup), the k-mer coverage (kcov), the user specified K-mer size (k), reads that contained errors (err), and the ploidy (p).

**Figure 3.**
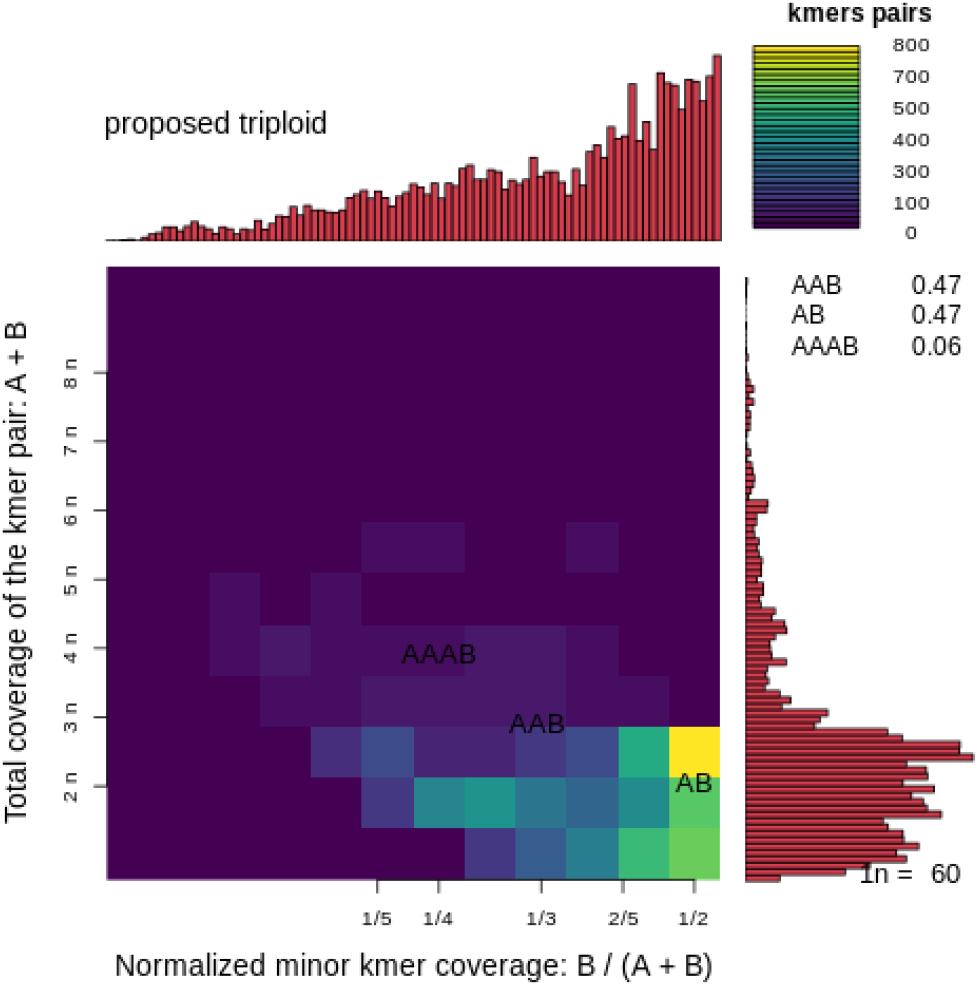
A smudgeplot providing the estimated ploidy level of *G. radlkoferi* genome. This figure describes the total coverage of the kmer pairs: A+B.

This was followed by the genome assembly with HiFiasm v0.25.0^14^ based on the HiFi reads. The assembly was then subjected to purging to remove haplotypic duplication and contig overlaps in the draft assembly using Purge_dups v1.2.6^15^. Thereafter, the genome was evaluated using the gfastats v 1.3.11 tool^16^. The genome assembly statistics obtained for *G*.*Radlkoferi* are compared with three plant species within the order of Geraniales in Table 1. Blobtoolkit and Blobtoolkit viewer v4.1.0^17^ were used to generate a snailplot to visually represent assembly statistics (Figure 5). Finally, BUSCO v5.8.0 was used in Galaxy with the Viridiplantae_odb10 lineage to assess the quality of the genome assembly.

**Table 1.**
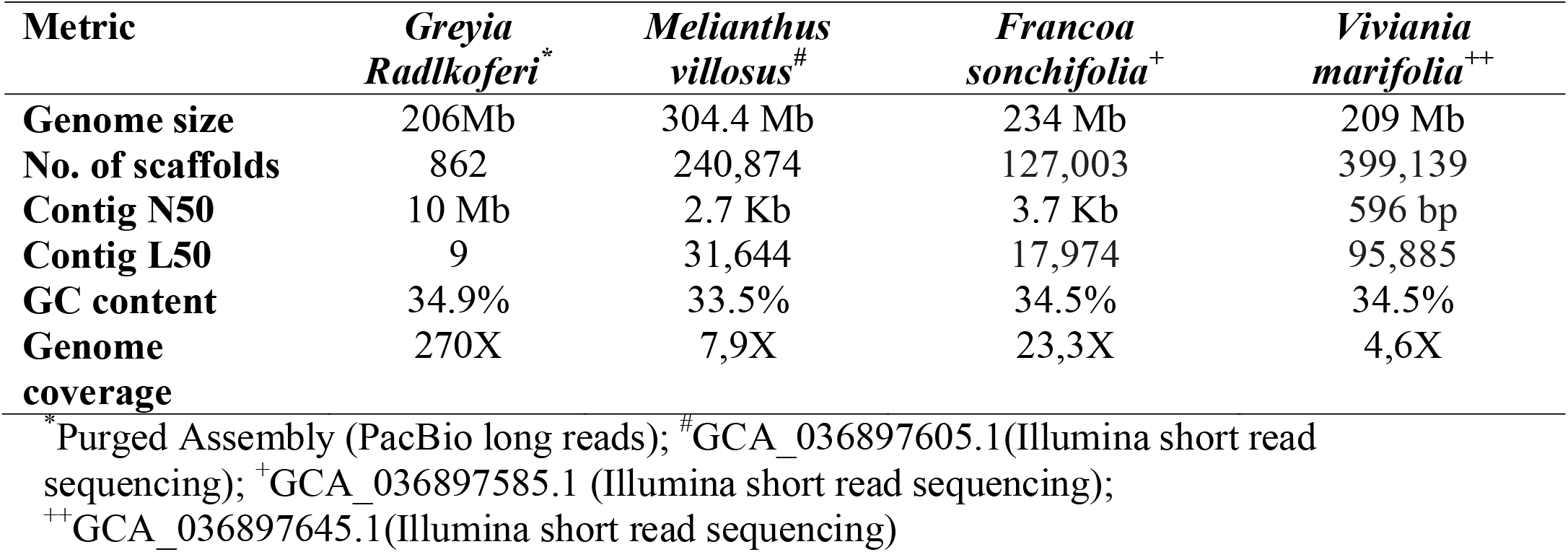
Assembly statistics of the purged Woolly bottlebrush, in comparison to other three plant species within the order of *Geraniales*.

### Genome quality and assessment

The genome size using genome scope for *G. radlkoferi* was estimated at 235 Mb through k-mer analysis at k = 17 (Figure 2). In the Smudgeplot (Figure 3), *G. radlkoferi* was determined to be triploid, with the following genotype frequencies AAB = 0,47 and AB = 0.47 and AAAB = 0.06. The BUSCO plot analysis indicated 98.4% completeness with 418 single-copy genes (Figure 4). The genome size after purging of duplicates was estimated at 206 Mb, with contig N50 of 10 Mb and contig L50 of 9.

**Figure 4.**
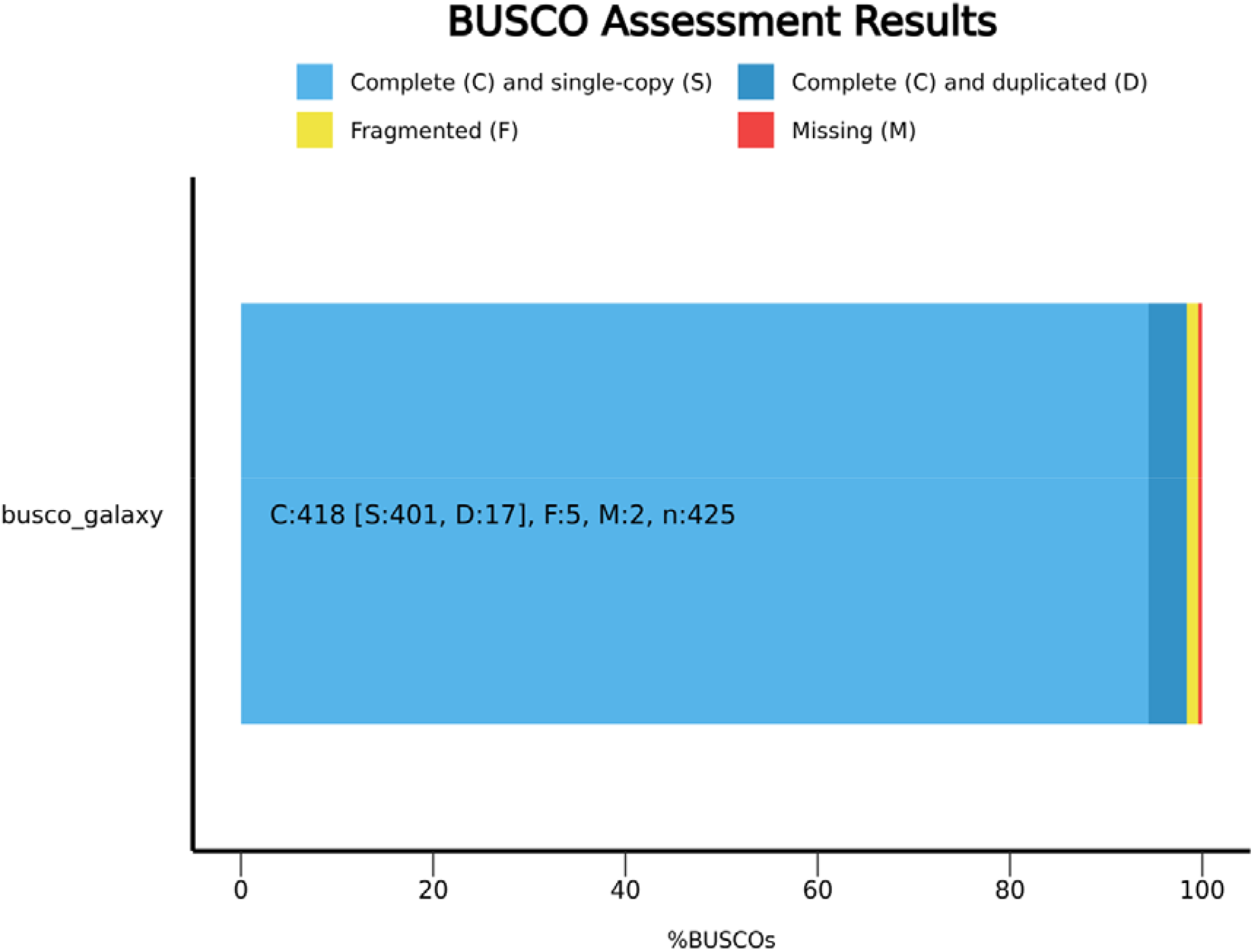
BUSCO plot of *G. radlkoferi* indicating the complete and single-copy genes of the assembled genome.

**Figure 5.**
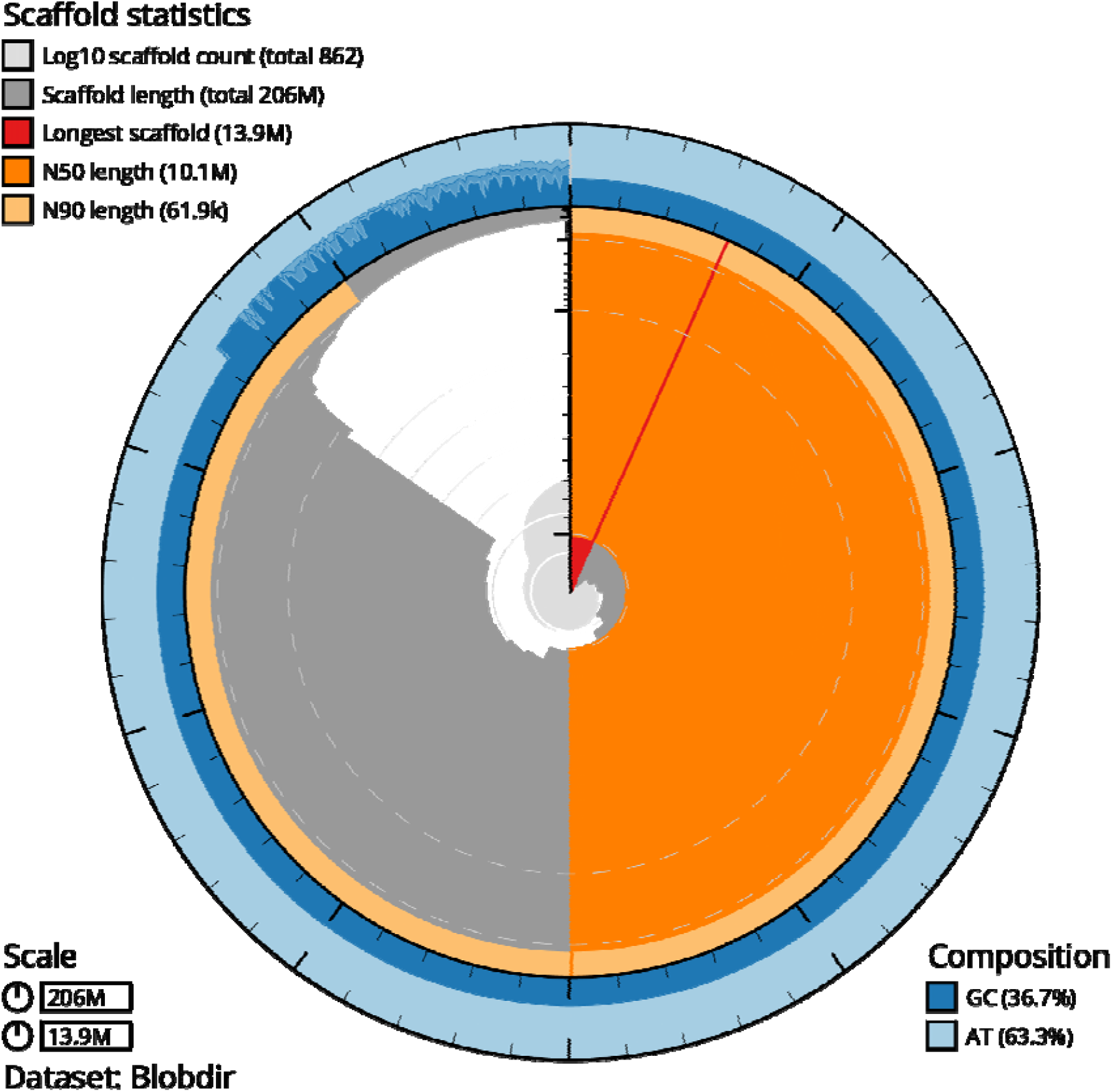
Blob figure depicting the scaffold statistics for the *Greyia radlkoferi*, providing the scaffold length and scaffold N50.

### Genome annotation

Repeat elements in the *G. radlkoferi* genome were found using RepeatModeler v.0.9^18^ and masked using RepeatMasker v4.1.5^19^ as the initial stage of genome annotation. RepeatModeler v0.9^18^, which includes tools such as RepeatScout v1.0.6, RECON v1.5.0, and TRF v4.09, was used to perform de novo repeat modelling to define *G. radlkoferi*-specific repeated elements. The Perl script “one_code_to_find_them_all”^20^ was further used to annotate the resulting repeats. Repeat analysis revealed that approximately 52,29% of the genome consists of repetitive elements. Retroelements are the most abundant among the repeats (7.56%). Among these, the most abundant are Long Terminal Repeats (LTR) elements, (6.99%), followed by Long Interspersed Nuclear Elements (LINEs) (0.43%). DNA transposons contributed a smaller proportion (1.65%), with contributions of hobo activators of 0.49%, with 12.06% comprising unclassified repeats which may represent species-specific or highly degenerate sequences. Other repeat classes include small RNAs (7.57%), simple repeats (1.69%), and low-complexity sequences (0.36%) (Table 2).

**Table 2.**
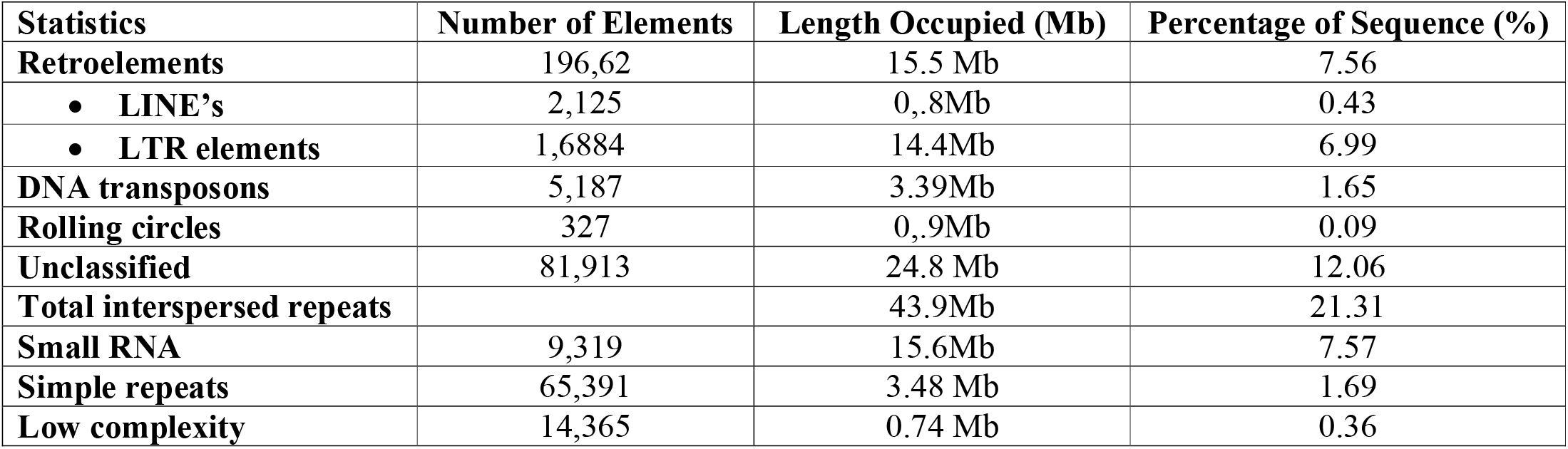
Predicted repeat element statistics for the *Greyia radlkoferi* genome generated through RepeatMasker.

| Statistics | Number of Elements | Length Occupied (Mb) | Percentage of Sequence (%) |
| --- | --- | --- | --- |
| **Retroelements** | 196,62 | 15.5 Mb | 7.56 |
| • LINE's | 2,125 | 0,8Mb | 0.43 |
| • LTR elements | 1,6884 | 14.4Mb | 6.99 |
| **DNA transposons** | 5,187 | 3.39Mb | 1.65 |
| **Rolling circles** | 327 | 0,9Mb | 0.09 |
| **Unclassified** | 81,913 | 24.8 Mb | 12.06 |
| **Total interspersed repeats** |  | 43.9Mb | 21.31 |
| **Small RNA** | 9,319 | 15.6Mb | 7.57 |
| **Simple repeats** | 65,391 | 3.48 Mb | 1.69 |
| **Low complexity** | 14,365 | 0.74 Mb | 0.36 |

The masked genome was annotated using Tiberius v1.1.8, a deep learning-based ab initio gene structure prediction tool. The results indicated the presence of 17,804 protein-coding genes, with an average exon length of 284.27 bp. The number of transcripts was 17,804, with an average gene length of 3,116.03 bp as shown in Table 3.

**Table 3.** Structural annotation statistics of the *Greyia radlkoferi*.

| Structural annotation | *Greyia radlkoferi* |
| --- | --- |
| **Number of protein-coding genes** | 17,804 |
| **Number of transcripts** | 13,845 |
| **Total number of exons** | 79,207 |
| **Total number of introns** | 61,403 |
| **Average gene length (bp)** | 3,116.03 bp |
| **Average exon length (bp)** | 284.27 bp |
| **Total CDS length (Mb)** | 22.5 Mb |
| **Total exon length (Mb)** | 22.5 Mb |
| **Total intron length (Mb)** | 32.9 Mb |

### Synteny

The synteny analysis between the *Greyia radlkoferi* and the *Francoa sanchofolia* (Figure 6) indicated a strong colinear relationship. Even though the genome size were different, about 75,6% of the Francoa genome aligned with the Woolly bottlebrush, suggesting that there are conserved chromosomal structure between these plant species.

**Figure 6.**
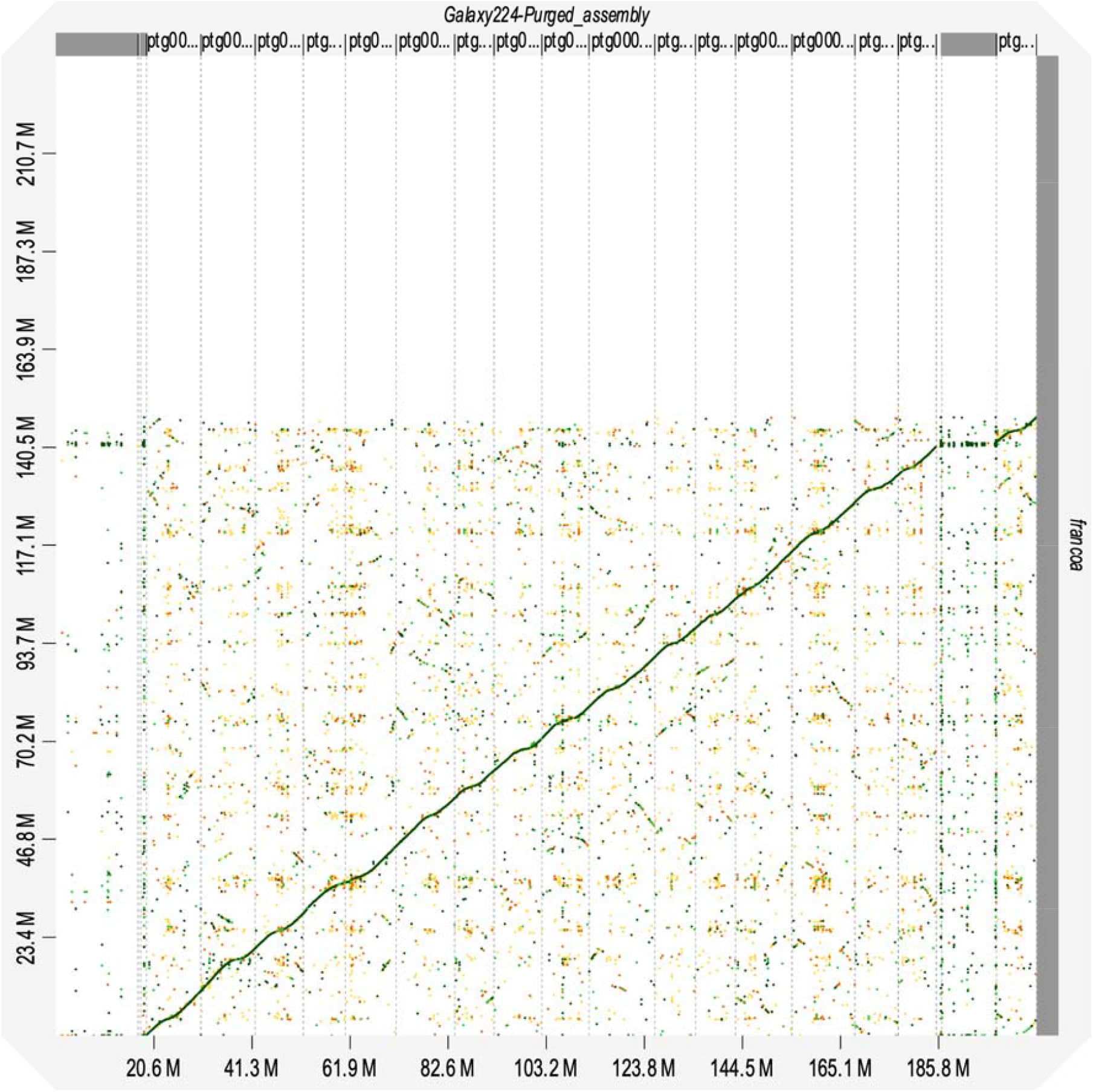
A collinearity plot from D-genies showing sequence similarity between assembled scaffolds of *Greyia radlkoferi* and *Francoa sancholia*. *The dot plot compares genomic coordinates between the reference (x-axis) and query (y-axis) genomes. Each point represents a homologous gene pair identified through sequence similarity. Green dots indicate collinear matches in the same orientation, while orange dots represent matches in reverse orientation (inversions). The prominent diagonal line reflects strong synteny and conserved gene order between the two genomes. Scattered off-diagonal points suggest rearrangements, duplications, or translocations. Chromosome or scaffold boundaries are indicated by dashed vertical lines, and axis scales (in megabases, Mb) denote genomic positions*.

### Orthogroups statistics

The total number of orthogroups identified across all the species was 33,770, with 88.1% of the genes assigned to orthogroups and 11.9% unassigned. A total of 88 single-copy orthologs were detected.

The comparative analysis indicated that 10368 orthogroups were shared between all the plant species analysed, which could indicate conserved plant functions among the species (Figure 7). Furthermore, there are 5961 orthogroups unique to *G*.*radlkoferi*, suggesting lineage-specific gene expansion that could be associated with the ability of the plant to adapt to different environments.

**Figure 7.**
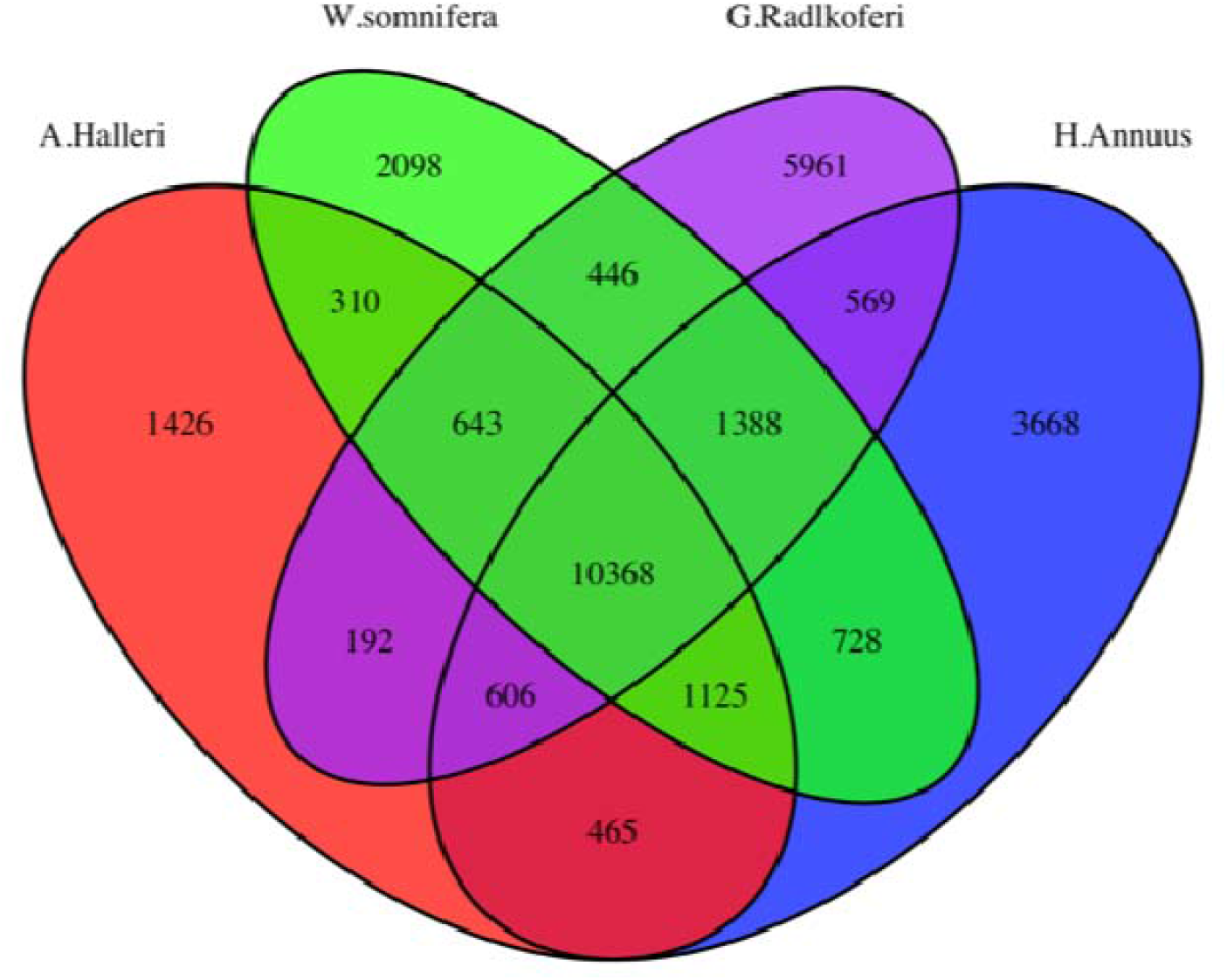
Venn diagram depicting the shared orthogroups between four different plant species.

### Phylogenetic analysis

For the phylogenetic analysis involving multiple sequence alignment of long-read sequencing data, protein sequences were required for the selected plant species. However, *Melianthus villosus, Francoa sonchifolia* and *Viviania marifolia* did not have protein sequences available, in public databases, and were therefore excluded from the analysis. *Vitus Vinifera, Withania somnifera, Helianthus annuus, Arabidopsis halleri*, and *Glycine max* were included in the phylogenetic analysis against *G. radlkoferi as* these species belong to the Eudicots and Rosids clades and have well-annotated protein datasets available in NCBI. To perform the analysis, Orthofinder v 3.1.0^21^ was used to identify orthologous gene groups across the selected species. The resulting phylogenetic tree illustrates the evolutionary relationships between *G. radlkoferi* and the selected plant species. According to the tree (Figure 8), *Withania somnifera* and *Helianthus Annuus* stems from a common node indicating closer genetic relatedness among these species. *G. radlkoferi* shares a common ancestor with *Arabidopsis helleri* and *Glycine max but* exhibit greater genetic divergence from these species, which may suggest an earlier evolutionary separation within this lineage.

**Figure 8.**
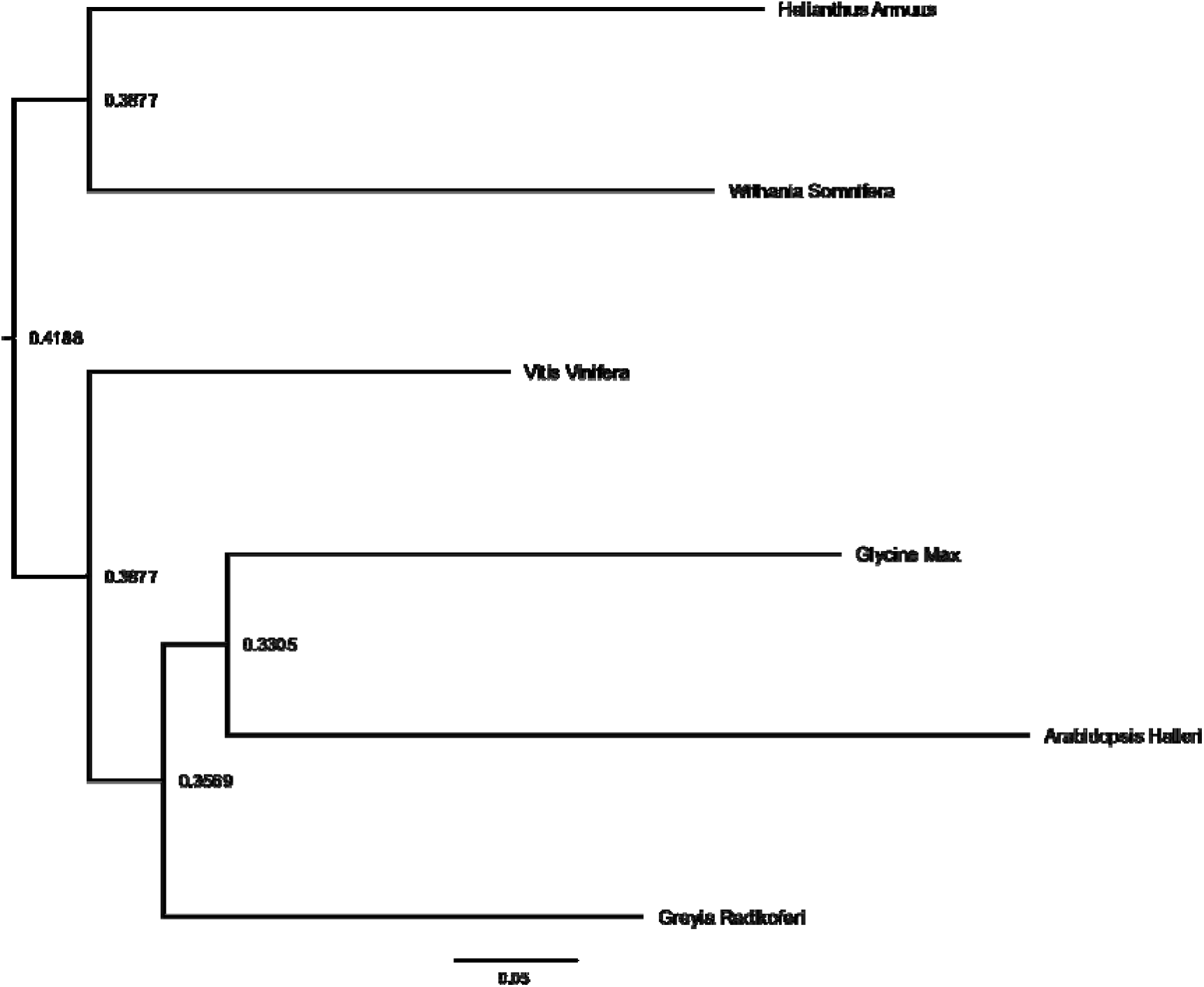
Phylogenetic tree showing evolutionary relationships among selected plant species from single-copy orthologs. The tree illustrates the inferred evolutionary distances between Vitis vinifera, Withania somnifera, Helianthus annuus, Greyia radlkoferi, Arabidopsis halleri, and Glycine max. Branch lengths are proportional to genetic distance, with numerical values along branches indicating the extent of divergence between nodes. The scale bar (0.1) represents the number of substitutions per site. Nodes represent common ancestors, and the topology reflects the relative relatedness among the species included in the analysis.

### Gene ontology analysis

The gene ontology (GO) terms for *G. Radlkoferi* were detected using the Interproscan version 5.59-91.0^22^ on Galaxy Europe platform^10^. Due to *G. Radlkoferi* being poorly annotated, gProfiler^23^ was used to convert the gene IDs to their corresponding annotated orthologs in *Arabidopsis Thaliana*, using the g:Convert function. Functional profiling for the gene IDs was done using the g:GOst function in gProfiler to obtain the Molecular Function (MF), Biological processes (BP), KEGG and Cellular Component (CC) pathways. For MF, six GO terms were identified, with protein binding being the most significantly enriched function. For BP, 46 significant GO terms were detected, with anatomical structure development being the most significant biological pathway. For CC, 14 GO terms were identified, with intracellular anatomical structure being highly significant (Figure 9). Genes associated with MF included *RPT5B, CR2, AB12, HAT*, while genes associated with BP included *A6, ELD1, TIM, COX11* and *FLC*. Furthermore, for the cellular component, *ATXR3, B160, CTF7*, and *ZFND3* genes were highlighted as significantly associated genes (See Figure S4). Histone acetyltransferase (*HA*T) is known to influence chromatin structure modification and assist in abiotic stress to drought, salinity and cold environments^24^. *CR2* was found to play a role in fungal resistance in white pine trees^25^.

**Figure 9.**
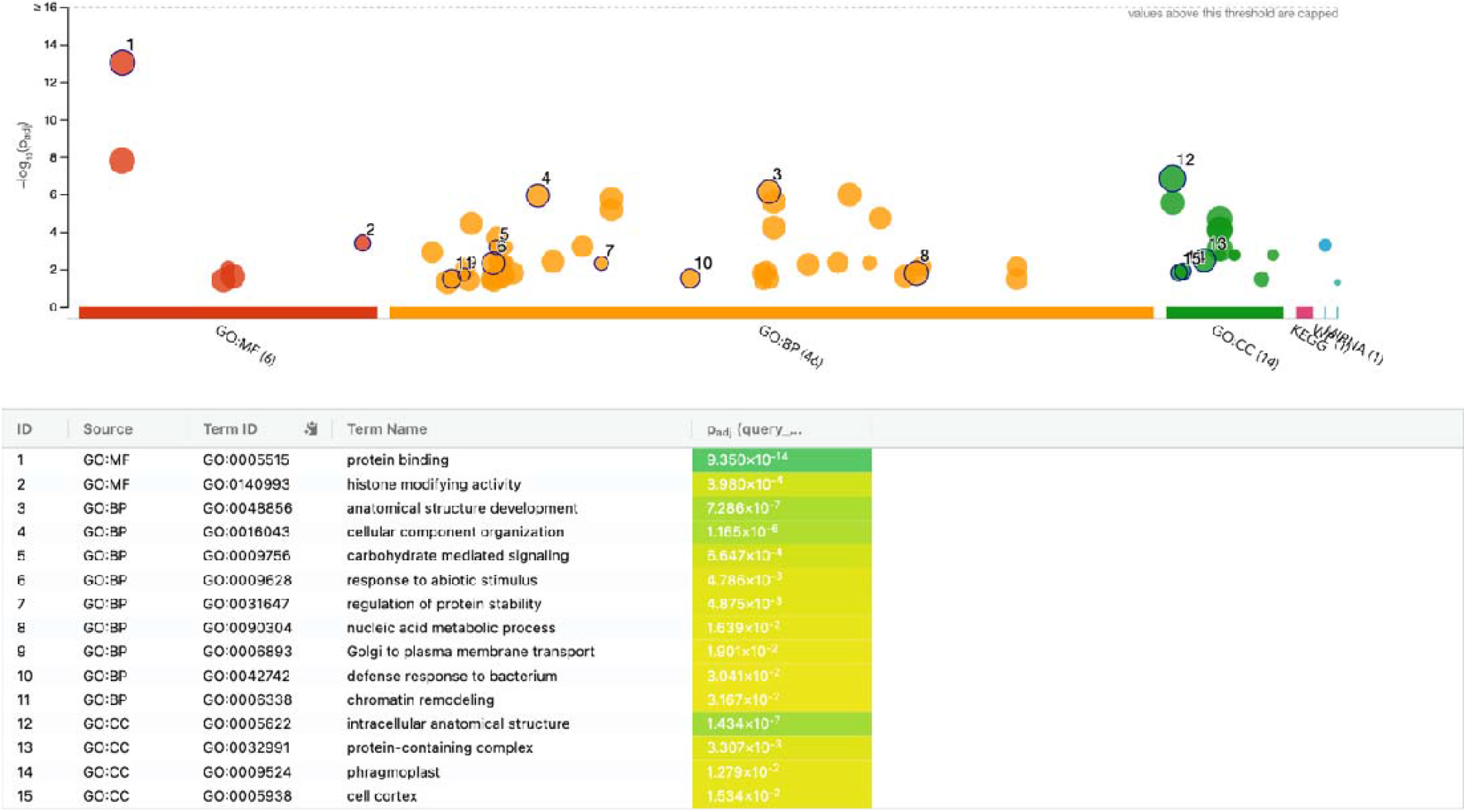
Gene ontology analysis for the molecular function, biological processes and cellular component using gProfiler for *Greyia Radlkoferi*.

This is the first draft genome of the *Greyia Radlkoferi* at 270X coverage. Even though it is not assembled at chromosomal level yet, it can be used as a reference genome for the other two plant species (*G. flanaganii* and *G*.*sutherlandii*) within the Greyia genus. This genome has been functionally annotated, and this data will be of value for future downstream analysis linking the regulation of genes that influence the expression and production of Greyia extracts utilised in the treatment of skin pigmentation in humans.

### Data Record

The raw sequencing data files are available on NCBI under the Bioproject PRJNA1227266, under the SRA section as two data files. The PacBio HiFi reads are in fasta format (SSR - PacBio_SMRT, Sequell II) in fasta format. Furthermore, the primary assembly https://doi.org/10.6084/m9.figshare.31812736 and alternate assembly files https://doi.org/10.6084/m9.figshare.31812871 in fasta format are available on Figshare. The genome annotation data is available on Figshare in GTF format.

## Technical Validation

### DNA sample quality

The High molecular weight DNA molecules were quantified using the Qubit 2.0 Fluorometer with the Qubit 1X dsDNA high sensitivity assay kit. DNA fragment length was determined using the Agilent Fragment Analyzer to confirm DNA concentration obtained from Qubit. DNA concentration value of 80ng/ul and fragment lengths on average of 15-20 bp were used for further library preparations. A minimum of 1ug DNA was used on the flow cell for Sequel IIe with 24-hour movie time to generate HiFi reads.

### Sequencing data assessment

The Phred score for all the sequencing data was above Q30, which is an indication that the base calling rate was more than 90% accurate. This is also provided in Table 1.

### Assessment of genome assembly quality

The BUSCO was used for the Benchmarking Universal Single-Copy Orthologs (BUSCO) and showed a high completeness of 98.4 % for Woolly bottlebrush with complete and single-copy (S). The error rate for both the primary and alternate assembly, as calculated through mercury was 1.68627e-06, with a QV value of 57 and completeness percentage of 99 % (Supplementary Table S1 and Figure S1).

## NCBI submission accession number

### Authors contributions

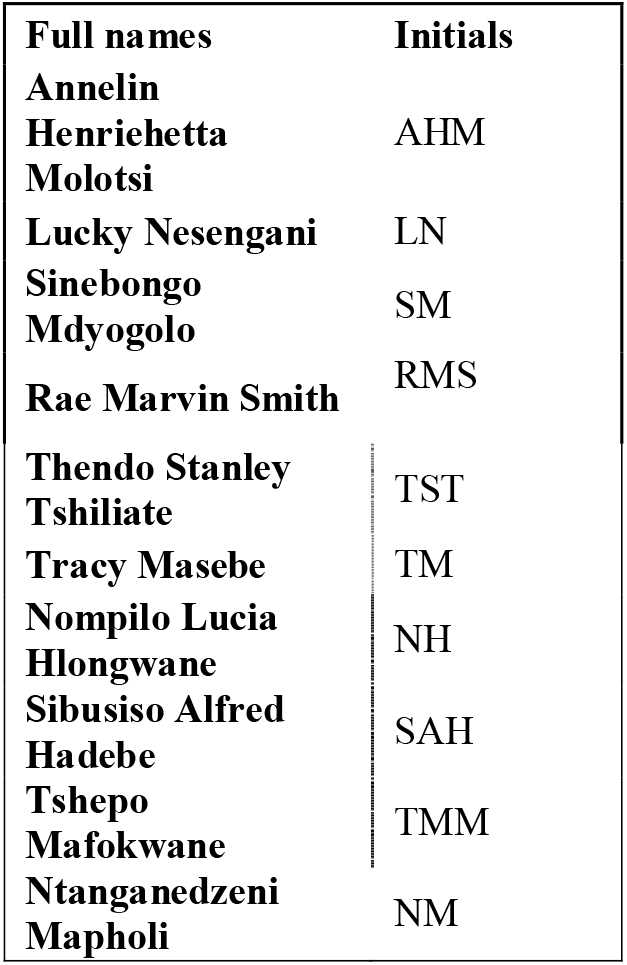

Conceptualization: AHM; Data Curation: AHM, LN, RMS; Formal Analysis: AHM, TST, NH; Funding Acquisition: NM; Investigation: AHM; Methodology: AHM, LN, SM, RMS; Project Administration: NM, TM; Resources: NM Software: AHM; Validation: LN, RMS, SM, TST, NM, TM; Visualisation: AHM, NH; Writing of the original draft: AHM; Reviewing and editing of the manuscript: AHM, LN, RMS, TST, TM, SM, SAH, NH,TMM.

## Data Availability

The completed genome assembly and raw data for the Woolly bottlebrush were submitted to the National Centre for Biotechnology Information (NCBI) under the Bioproject PRJNA1227266. With the following SRA accession numbers: SSR

## Code availability

All the analyses done in the current study were processed by employing the VGP pipeline, and RepeatModeler and RepeatMasker for the identification of repetitive elements and masking of these elements and Tiberius for annotation. All the commands and pipelines were executed following the manual and protocols of the corresponding bioinformatics software. In all the workflows, unless mentioned and where necessary, we used the default parameters.

## Acknowledgements

This project is supported by the University of South Africa, Africa-nuanced sustainable development goals research support programme. The samples for this project and the technical support were provided by the Africa BioGenome Project. A special thanks to Mapholi labs for providing the resources and managing the project.

